# Is constitutive red-shift an advantage for oxygenic photosynthesis under M-dwarf starlight? Insights from *Acaryochloris marina* sp. str. Moss Beach

**DOI:** 10.64898/2026.04.21.719884

**Authors:** Elisabetta Liistro, Beatrice Boccia, Mary Niki Parenteau, Nancy Y. Kiang, Nicoletta La Rocca

## Abstract

In the next years, several space missions will search for evidence of life on exoplanets, focusing on robust biosignatures associated with oxygenic photosynthesis, including atmospheric oxygen accumulation and the Vegetation Red-Edge in surface reflectance spectra. Many potentially habitable rocky exoplanets orbit M-dwarf stars, whose spectral energy distribution may challenge oxygenic photosynthesis. Differently from the Sun, M-dwarf stars emit predominantly far-red (700– 750 nm) and infrared (750–1000 nm) light, and relatively little visible (400–700 nm) radiation, which constitutes photosynthetically active radiation. Some organisms have been found to photosynthesize under such spectrum but less efficiently than under solar light, as their photosynthetic apparatus evolved to harvest visible light emitted by the Sun. Around M-dwarfs, such different irradiation might have selected adaptations optimized for harvesting far-red / infra-red light. On Earth, similar selection can be found in *Acaryochloris marina* strains, constitutively presenting high chlorophyll *d* content in photosystem II & I, with *in vivo* absorption peaks beyond 700 nm. Here we tested the Moss Beach strain under a simulated M-dwarf spectrum and a simulated primeval atmosphere – anoxic and enriched in carbon dioxide. Results underline how this permanently red-shifted photosynthetic apparatus does not require acclimation to the stellar spectrum and enables for a strong growth and oxygen production, higher than under simulated solar light. Moreover, cells’ reflectance spectrum highlights a shift of the canonical red-edge toward longer wavelengths, resulting in a *Chl* d-*near-infrared edge*, suggesting a similar metabolism on exoplanets orbiting M-dwarfs could successfully produce both a gaseous biosignature and a characteristic surface biosignature.

**Graphical abstract:** 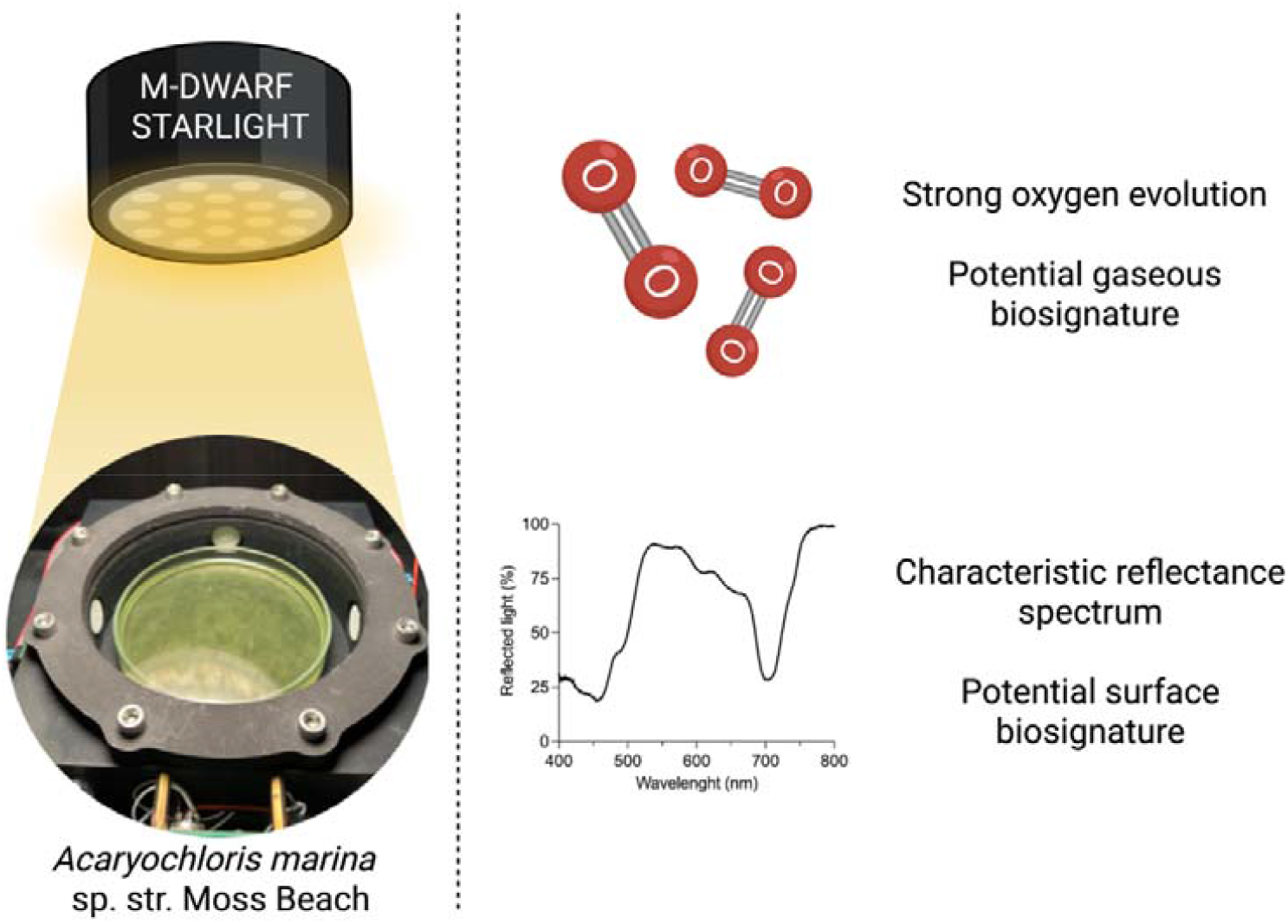

Created in BioRender. Liistro, E. (2026) https://BioRender.com/j2de4ay

## 1. Introduction

In the last decades, space missions such as Kepler, TESS (Transiting Exoplanet Survey Satellite, NASA) and GAIA (Global Astrometric Interferometer for Astrophysiscs, ESA), identified a large number of exoplanets, over 6000 at the time of writing^1^. Some of these planets have been recognized as potentially habitable as their features and of their host star(s) may support the presence and evolution of life (Kasting et al., 1993; Kaltenegger, 2017; Méndez et al., 2021). Among these, rocky Earth-like exoplanets were found to be common around M-dwarf stars (Kopparapu et al., 2013; Hsu et al., 2020). These stars, also referred to as M stars or Red-dwarfs, are considered good targets for the search for life as not only they frequently host rocky planets but they also have a lifespan to be long enough to allow for a biosphere to evolve (Shields et al., 2016). Moreover, their strong early stellar activity is considered to decline to solar levels within approximately 3 billion years, thus it does not necessarily preclude habitability (Gale & Wandel, 2016; O’Malley-James & Kaltenegger, 2017; Schwieterman et al., 2018).

In the frame of searching for life on planets orbiting M-dwarf stars, *in situ* detection is not feasible, as the closest M-dwarf, Proxima Centauri b, is distant 4.2 light-years from us (Anglada-Escudé et al., 2016). Life detection must thus rely on remote sensing of *biosignatures*, patterns or molecules of unequivocal biological origin (Schwieterman et al., 2018; Claudi & Alei, 2019). Missions such as ARIEL (Atmospheric Remote-Sensing Infrared Exoplanet Large-survey, ESA) and HWO (Habitable Worlds Observatory, NASA) will be launched in the coming years with the purpose of characterizing exoplanets atmospheres, looking for biosignatures (Feinberg et al., 2026; Tinetti et al., 2018). One of the strongest remotely detectable biosignatures is the presence of oxygen, O_2_, and its byproduct ozone, O_3_, in a planet’s atmosphere (Schwieterman et al., 2018; Cavalazzi & Westall, 2019; Claudi & Alei, 2019). Since on Earth the abundance of these molecules is caused by oxygenic photosynthesis, the metabolic process operated by cyanobacteria, algae and plants (Cavalazzi & Westall, 2019), scientists have been discussing on the feasibility of this metabolisms around M-dwarf stars (Tinetti et al., 2006; Kiang et al., 2007a; Kiang et al., 2007b).

M-dwarfs are cold and faint stars that emit mainly far-red (FR, 700–750 nm) and infra-red (IR, 750–1000 nm) light, and very low visible (VIS, 400–700 nm), in contrast to G stars, like the Sun (Chabrier et al., 1996; Pecaut & Mamajek, 2013), whose emission is stronger in the visible. This emission spectrum challenges the possibility of oxygenic photosynthesis as this process on Earth relies mainly on VIS light, also referred to as *photosynthetically active radiation*, PAR. Nonetheless, models indicate that the small amount of VIS light reaching a planet orbiting an M-dwarf star could still allow for oxygenic photosynthesis to work efficiently, with a productivity estimated to be up to the 22% of that on Earth (Gale & Wandel, 2016; Ritchie et al., 2018; Gray et al., 2025). If photosynthetic adaptations such as three or four photosystems in series have evolved around M-dwarf stars, the productivity of oxygenic photosynthesis has been estimated to potentially increase to over the 75% of Earth’s one (Kiang et al., 2007b). Additionally, oxygenic photosynthetic organisms generate another biosignature among the most studied: the Vegetation Red-Edge, VRE. VRE is a sharp increase in reflectance around 700 nm, as a consequence of the contrast among the absorption of visible wavelengths, due to chlorophyll molecules, and the scattering of near-infra-red wavelengths due to cell structures (Sagan et al., 1993; Turnbull et al., 2006; O’Malley-James & Kaltenegger, 2018; Schwieterman et al., 2018). It was proposed that, around an M-dwarf, this feature of (eventual) oxygenic phototrophs could be altered because of the need to harvest longer wavelengths (Wolstencroft & Raven, 2002). Due to the presence of FR or near-infra-red (NIR) harvesting pigments, the VRE could be shifted and become a “NIR” edge (Tinetti et al., 2006; Kiang et al., 2007a).

Recently, it was experimentally demonstrated that oxygenic photosynthesis and growth is possible under a simulated M-dwarf spectrum in several Earth organisms, with different efficiencies (Battistuzzi et al., 2023a; Battistuzzi et al., 2023b; Liistro et al., 2024). A deep investigation was carried out for cyanobacteria, as in the last decades some strains were found capable of photosynthetizing not only with visible light but also with solely far-red light via the Far-Red Light Photoacclimation (Gan et al., 2014). In these species, the FaRLiP response is enabled by the presence of a specific gene cluster expressed in FR light which leads to the synthesis of red-shifted pigments, chlorophyll *d* (Chl *d*), chlorophyll *f* (Chl *f*) and the accessory pigment far-red allophycocyanin (FR-APC), together with paralog subunits of photosystems that incorporate the pigments. With this remodeling of the photosynthetic apparatus, organisms present a shoulder of absorption in the far-red and fluorescence emission shifted to longer wavelengths (Gan et al., 2014; Ho et al., 2017).

*Chlorogleopsis fritschii* PCC6912 acclimated to the simulated M-dwarf spectrum showed the activation of FaRLiP, leading to the use of both the visible and FR light available in the stellar spectrum (Battistuzzi et al., 2023a).

These results are extremely important to understand the limits of oxygenic photosynthesis around M-dwarfs, however, they all consider organisms evolved for photosynthesizing with visible light and that canonically harbor VIS photosystems, with some exception of strains that at most can acclimate to increase their far-red absorption capacity through uphill energy transfer (Elias et al., 2024). It was hypothesized that the eventual evolution of oxygenic phototrophs under the irradiance of M-dwarfs could have selected particular adaptations such as red-shifted pigments, modified antennae, or a higher number of photosystems working in series (Kiang et al., 2007a; Kiang et al., 2007b; Takizawa et al., 2017; Chitnavis et al., 2024).

Curiously, on Earth we have examples of organisms, adapted to FR-enriched light niches, with a unique photosynthetic apparatus that has constitutive and severe red-shifted absorbance, namely in the cyanobacterial strains of the species *Acaryochloris marina* (Miyashita et al., 1996; Larkum & Kühl, 2005). *A. marina* strains are characterized by the permanent large presence of Chl *d* in Photosystem I & II, accounting for more than 90% of the total chlorophyll content, and participating both in light harvesting and charge separation, with maximum absorption at 740 nm (Chen et al., 2002; Badshah et al., 2017). It is also interesting to notice how ancestral the divergence of this genus is in the evolution of oxygenic photosynthesis: the split of the *Acaryochloris* genus from the cyanobacterial species tree was found to precede the appearance of the FaRLiP phenotype, dated in the Paleoproterozoic (Antonaru et al., 2025).

When considering the eventual evolution of oxygenic photosynthesis around M-dwarfs, *A. marina* is an excellent example of the evolution of a photosynthetic apparatus that mainly harvests FR wavelengths.

In this work, we evaluated the response of *Acaryochloris marina* sp. str. Moss Beach to simulated planetary conditions, comprehensive of an M-dwarf simulated spectrum and an anoxic atmosphere enriched in CO_2_.

*A. marina* sp. str. Moss Beach was found as an epiphyte of a brown macroalga, in the rocky intertidal zone of the J.V. Fitzgerald Marine Reserve, California, and the Chl *d* in its photosynthetic apparatus can reach up to 99% of the total chlorophyll content (Kiang et al., 2022). Due to its Chl *d-*bearing photosystems, the *in vivo* Q_y_ absorption peak of *A. marina* sp. str. Moss Beach is at 705 nm, with its tail arriving at 750 nm. This strain lacks the accessory pigments phycobiliproteins, hence the light harvesting is only operated with chlorophylls and marginally with carotenoids, similarly to other strains of *A. marina*, as *A. marina* CCMEE 5410 and *A. marina* HICR111A (Chan et al., 2007; Mohr et al., 2010; Kiang et al., 2022).

As this strain is of recent discovery, we first identified the medium, temperature and light intensity suitable for the simulations. We then assessed whether *A. marina* sp. str. Moss Beach performed a physiological acclimation to a simulated M-dwarf spectrum, comparing the response with cells exposed to solely FR light and solar light. Finally we proceeded by investigating its performances in simulated planetary conditions combining irradiation with a simulated anoxic primeval atmosphere.

## 2. Materials and methods

### 2.1 Cultivation conditions for preliminary experiments

We obtained the initial inoculum of *Acaryochloris marina* sp. str. Moss Beach, isolated at Moss Beach, in the J.V. Fitzgerald Marine Reserve, California from co-authors Kiang and Parenteau (Kiang et al., 2022). The strain was maintained in liquid MBG11 medium at 25°C under 15 µmol of photons m^-2^s^-1^ of continuous white fluorescent light (L36W-840, OSRAM).

For the experiment testing different growth media and temperature, 50 mL inocula with initial optical density at 750 nm, OD_750_, of 0.2 were grown either in MBG11 medium or in ASNIII medium, a common medium for marine cyanobacteria. Cultures were exposed to 15 µmol of photons m^-2^s^-1^ of continuous white light (TRUE-LIGHT 18T8, Lightfull). For each tested medium, four biological replicates were grown at 20°C, at 25°C or at 30°C.

For the experiment testing light intensities, cultures with initial OD_750_ of 0.2 were grown at 25°C in liquid MBG11 in a multi-cultivator ((Multi-Cultivator MC 1000-OD, Photon Systems Instruments, Czech Republic). Three biological replicates were exposed to 10, 20, 40, 60, 80, 100 and 150 µmol of photons m^-2^s^-1^ of continuous cold white light provided by built-in LEDs in the multi-cultivator.

### 2.2 Cultivation conditions for planetary simulations

Planetary conditions in terms of irradiance and atmosphere were simulated with a custom-made set up extensively described in Battistuzzi et al., 2020, 2023a and Claudi et al., 2021. Briefly, the set-up is comprehensive of three light sources and three growth chambers.

The light source for planetary simulation is a custom-made *starlight simulator*, SLS, which for this work was used to reproduce the irradiation of an M-dwarf star (M7) in the range 350 nm – 850 nm (Figure S1). For comparison, cultures were exposed also to a simulated solar light (SOL) and a monochromatic far-red light (FR). SOL light was provided by a custom-made visible-infra-red light simulator with 4 independently manageable led channels in the range 365-800 nm. Channel 1 controls leds at 365 nm, 385 nm, 405 nm, 425 nm and 500 nm; channel 2 controls SUNLIKE 5000K and 690 nm leds; channel 3 (far-red) controls leds at 730 and channel 4 (infra-red) controls leds at 750 nm and 780 nm. Channel intensity was regulated in order to accurately reproduce the Sun emission spectrum (Figure S1). FR light peaked at 730 nm was provided with the illuminator described in Battistuzzi et al., 2023 (Figure S1a). For all light conditions, M7, SOL, FR, the total light intensity was 20 µmol of photons m^-2^s^-1^, resulting respectively in 7.38, 16.92 and 1.47 µmol of photons m^-2^s^-1^ of visible light and 12.62, 3.08 and 18.53 µmol of photons m^-2^s^-1^ of far-red light. A more detailed repartition of light in the different wavebands is available in Figure S1c.

For experiments investigating acclimation to the M7 starlight, 50 mL of refreshed cultures at OD_750_ 0.2 were transferred from white light under M7, SOL and FR light. Growth and *in vivo* absorption were monitored periodically for 21 days, after which cultures were examined for physiological changes indicating the display of eventual acclimation strategies.

For planetary simulations, where an anoxic primeval atmosphere was combined with stellar irradiation, *atmosphere simulating chambers*, ASCs, were used. ASCs are growth chambers that allow cultivation of organisms underneath chosen light spectral irradiances – in this study M7, SOL and FR (Figure S1b). The ASCs can be inflated with a desired gas composition. Inside the ASCs are one sensor for O_2_ concentration (LOX-O2-S, SST sensing) and two sensors for CO_2_ concentration (CO2M-20 and CO2M-100, SST sensing), which enable to monitor in real time and register the gas consumption and release during the experiment. In this work, the initial atmosphere inside the ASCs was composed of 95% nitrogen, N_2_, and 5% CO_2_, simulating an anoxic primeval planetary atmosphere. Raw data of gas concentrations in ppm registered during the experiments were converted in µmol through Matlab R2018b (MathWorks) by applying the law of ideal gasses. Experiments with ASCs were run in parallel under M7, SOL and FR for four biological replicates.

### 2.3 Growth assessment

Growth of *A. marina* sp. str. Moss Beach was assessed through evaluation of optical density at 750 nm, OD_750_, measured with a spectrophotometer (Cary100 UV-VIS, Agilent) against a blank (culture medium). Maximum growth rate µ was calculated on the steepest section of growth curves as follows:

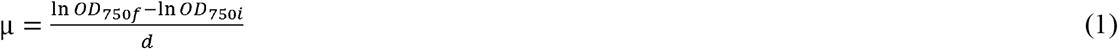

were *OD*_*750f*_ is the OD_750_ value at the end of the section, *OD*_*750i*_ is the value at the beginning of the section and *d* is the number of days passed between the two points.

### 2.4 In vivo *absorption, reflectance and fluorescence*

To record *in vivo* absorption spectra of cells, samples were centrifuged at 1400 g for 10 min at room temperature and the pellet was homogenized with a pestle. 600 µL of fresh medium was added to solubilize the pellet and samples were analyzed with a Cary100 UV-VIS spectrophotometer (Agilent) inside quartz cuvettes. The opaque side of the cuvette was exposed to the light ray to correct the scattering (Gan et al., 2014; Amesz et al., 1961; Shibata, 1959).

*In vivo* reflectance of cells was recorded with the set-up described in Liistro et al., 2026. Briefly, a halogen lamp (Halostar Starlite 12V 20W G4, Osram) was used to irradiate 5 mL of liquid cultures with OD_750_ of 5 in a petri dish and reflectance was collected in the range 400–1000 nm with an optical fiber connected to a spectrometer (Flame, OceanOptics) via the SpectraSuite software (OceanOptics). Aquired spectra were normalized against a white reference. To have control reflectance spectra for the one of *A. marina* sp. str. Moss Beach acclimated to the M7 simulated starlight, this analysis was carried out also on cultures of *Synechocystis* sp. PCC6803 and *Chlorogleopsis fritschii* PCC6912 acclimated to the same simulated stellar spectra. Information on the acclimation that these two strains perform under this condition can be found in Battistuzzi et al., 2023a, 2023c.

Low temperature (77K) *in vivo* fluorescence emission spectra were recorded with a Cary Eclipse Fluorescence Spectrometer (Agilent). 2 ml of culture were centrifuged at 1500 g for 10 min at 4 °C and the pellet resuspended in 1 ml of glycerol 60% (w/v) and 10 mM Hepes pH 7.5. Samples were frozen in liquid nitrogen and kept in dark at -80 °C until analysis. For the analysis, samples were excited at 440 nm and emission spectra were recorded between 600 nm and 900 nm. Low temperature (77 K, −196.15 °C) during recording of emission spectra was maintained with liquid nitrogen.

### 2.5 Pigment content

For the extraction of lipophilic pigments, samples were centrifuged for 10 min at 17500 g at 4 °C and to the pellet were added an equal volume of 150-212 µm acid washed glass beads (SIGMA) and 50 µl of HPLC grade methanol 100%. Three cycles of cell rupture were executed with a Bullet Blender Storm Pro Homogenizer (Next Advance) for 15 s at maximum speed alternated with 30 s in ice. 150 µl of methanol were added to the lysate and a last cycle of rupture was executed. Samples were centrifuged at 20000 g for 10 min at 4 °C and the supernatant, containing extracted pigments, was transferred to a new tube and kept in ice. 200 µl of methanol were added to the pellet and, after vortexing for resuspension, another centrifugation was performed. The supernatant was pooled with the previously obtained extract. These last steps were repeated until the supernatant appeared transparent, indicating full extraction of pigments. Samples were kept in dark at -20 °C until analysis. Pigment extracts were analyzed with a Cary100 UV-VIS spectrophotometer (Agilent). Concentration of chlorophyll *a* and chlorophyll *d* were obtained with the following equations (Ritchie, 2006):

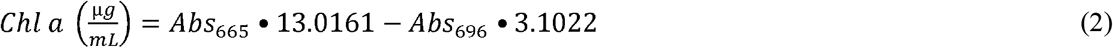

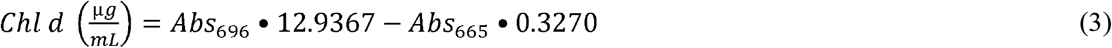

To evaluate the relative content of chlorophyll and carotenoids, RP-HPLC analyses were conducted on pigment extracts. Analyses were performed with an Agilent 1100 series LC with a Lichrospher 100 RP column (250 mm length, 4 mm diameter) containing 5 μm silica particles coated by C-18 atom chains (Merck) as the stationary phase. The mobile phase consisted in buffer A (methanol:acetonitrile:water in ratio 42:33:25) and buffer B (methanol:acetonitrile:ethyl acetate in ratio 50:20:30). Buffer A and B were eluted in the column according to the protocol optimized by Gan and collaborators (Gan et al., 2014). The relative content was assessed by integrating the area of each peak in chromatograms at 440 nm for four biological replicates.

### 2.6 Confocal imaging

*A. marina* sp. str. Moss Beach cells were imaged with a confocal microscope (STELLARIS 8, Leica) to check for eventual changes in dimension, morphology and chlorophyll localization. Cells were excited by a laser diode with emission at 405 nm and imaged with two different channels to separate the fluorescence signal from chlorophyll *a* (672–689 nm) and chlorophyll *d* (705–739 nm).

### 2.7 Statistical analyses

Statistical analyses, ANOVA analysis (one-way or two-way ANOVA depending on the dataset) and Tukey’s or Šídák’s multiple comparison tests, were performed with the software GraphPad Prism v10.1.0 (GraphPad).

## 3. Results

### 3.1 *Responses to nutrients, temperatures and light intensities in* A. marina *sp. str. Moss Beach*

Since *Acaryochloris marina* sp. str. Moss Beach is of recent discovery, little information is available about the impact of environmental parameters on its growth. Prior to the planetary simulation, we performed preliminary experiments investigating how different temperatures, media and light intensities affected the growth rate of this strain. Being originally developed for experiments on cyanobacteria, the simulator currently has the lower limit of operational temperature set at 25°C, which is slightly higher than the environmental temperature at which *A. marina* sp. str. Moss Beach was isolated, that is around 19°C. To understand whether this difference could have a non-negligible impact on the growth and performances of cells during the simulation, we followed their growth at three different temperatures: 20°C, reproducing the natural environment, 25°C, the target temperature, and 30°C, the temperature at which many cyanobacteria strains are maintained and at which we performed planetary simulations on other strains in the past (Liistro et al., 2024; Battistuzzi et al., 2023a).

Concurrently, we evaluated the growth in two different saline media, Marine BG11 (MBG11) and artificial seawater (ASNIII). MBG11, rich in iron, is the medium suggested for *A. marina*, but the presence of small agglomerates hinders the maintenance of full axenicity in the cultures. ASNIII, on the other hand, is a common medium for marine strains and facilitates the maintenance of clean cultures but has a lower salinity and a lower iron content.

Cultures were exposed to these 6 different conditions – 20°C, 25°C, 30°C in both MBG11 and ASNIII – under continuous irradiation of 15 µmol of photons m^-2^s^-1^ of white light. Growth was followed via monitoring of OD_750_ for 16 days (Figure 1). Considering the maximum growth rate registered during the experiment, the only temperature that was significatively affecting cell growth was 30°C, in both MBG11 and ASNIII. Even though final values of OD_750_ were higher in cells grown at 20°C than at 25°C, growth rates registered in the exponential phase do not differ significatively. Concerning the media, ASNIII supports a faster initial growth, although maximum growth rates at the same temperature are not significatively different from those promoted by MBG11. Nonetheless, after 7 days for cells at 25°C, and 12 days for cells at 20°C, a stop in the growth was encountered, suggesting that scarcity of essential nutrients in the ASNIII medium was reached in a shorter time than in MBG11, where OD_750_ keeps gradually increasing.

**Figure 1.**
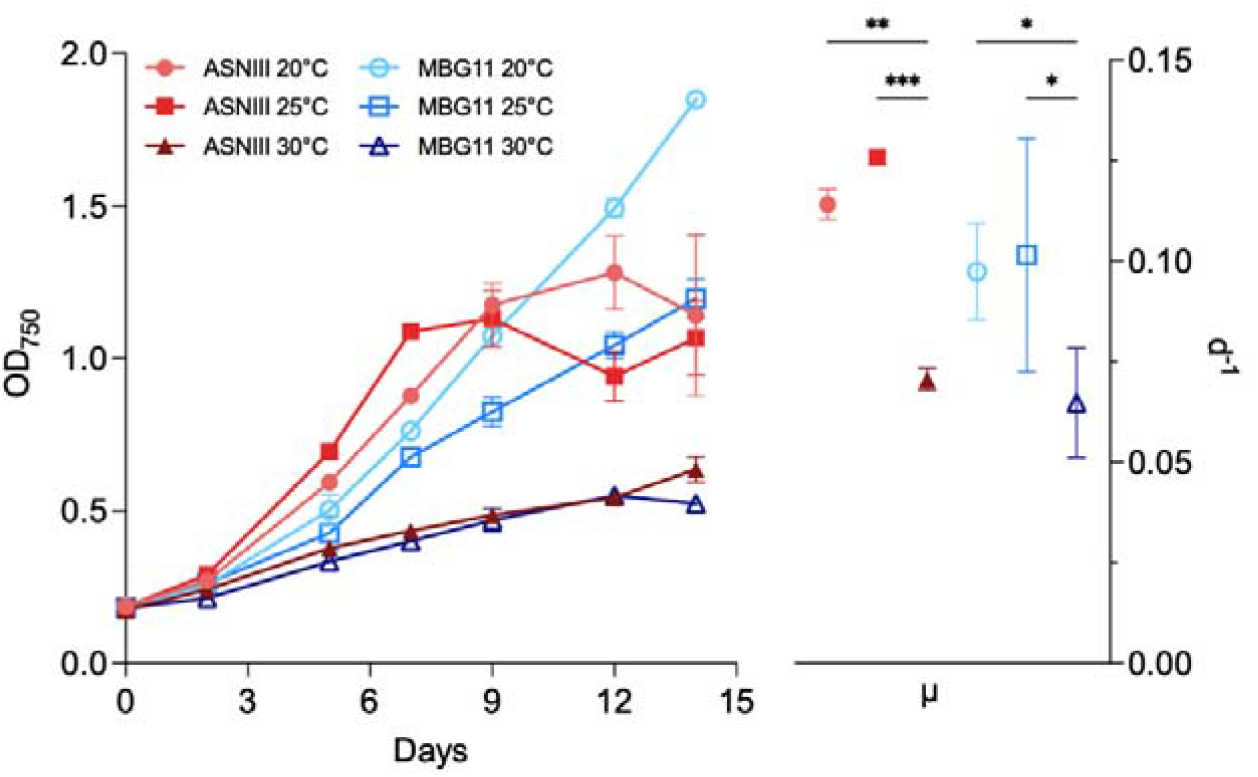
Growth of *Acaryochloris marina* sp. str. Moss Beach in different media and at different temperatures, in terms of values of OD_750_ and of growth rate (µ, d^-1^). Cells were grown in Marine BG11 (MBG11, blue shades) and in artificial seawater (ASNIII, red shades). For each medium, three temperatures were tested: 20°C (round), 25°C (square) and 30°C (triangle). OD_750_ was registered every 2-3 days (left), and, from OD_750_ values, the maximum growth rate (µ) for each condition was calculated (right). Data are presented as the mean of four biological replicates, for values of µ one-way ANOVA was employed, ***: p-val <0.0005, **: p-val <0.005, *: p-val <0.05. ASNIII: artificial seawater medium; MBG11: Marine BG11 medium.

To assess whether the composition and organization of photosynthetic apparatus was affected by the tested condition, the diagnostic trait of Chl *a* and *d* content was analyzed. No significant differences were registered in any of the conditions in terms of µg of pigment per OD_750_ (Figure 2), indicating cells maintain stable content of both Chl *a* and *d*. A slight difference was in cells grown at 30°C, which showed a higher percentage of Chl *d* both in ASNIII (98.23%) and in MBG11 (97.15%) than what registered in cells grown at 20°C and 25°C (around 94.50%).

**Figure 2.**
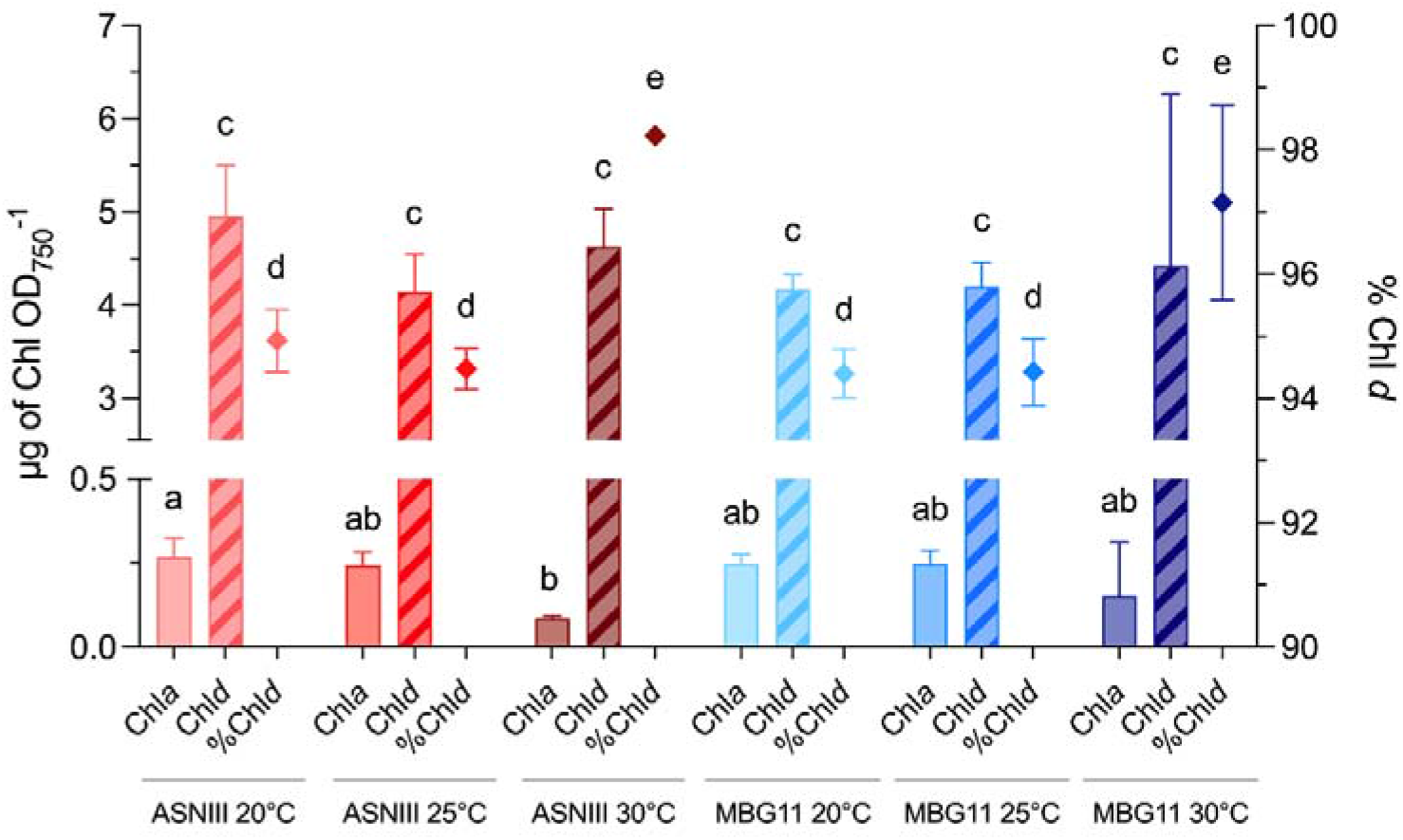
Pigment composition in *Acaryochloris marina* sp. str. Moss Beach grown in different media, ASNIII (red shades) and MBG11 (blue shades) and at different temperatures, 20°C (light shades), 25°C (normal shades), 30°C (dark shades). Chlorophyll *a* and *d* content are expressed as µg of pigment per OD_750_, percentage of chlorophyll *d* on total chlorophyll content (rhombus); statistical significance was assessed with one-way ANOVA on at least three biological replicates. ASNIII: artificial seawater medium; MBG11: Marine BG11 medium; Chl *a*: chlorophyll *a*; Chl *d*: chlorophyll *d*.

Considering the results of this preliminary screening, for further experiments cells were maintained in MBG11, which provides the appropriate amount of nutrients, at our target temperature of 25°C, since neither the growth nor the photosynthetic apparatus seem to be significatively affected.

With these parameters, experiments to identify the optimal light intensity for the acclimation to the M7 starlight were carried out. Indeed, prior to the simulation of planetary conditions, a step of acclimation to the stellar spectrum is needed, to understand whether a light with different spectral qualities triggers any change in the photosynthetic apparatus ascribable to any acclimation strategy. Acclimations may require time – days or weeks – to be implemented and detectable, and an evaluation of the optimal light intensity for the acclimation period is needed. In this case, the aim was to find a non-saturating light intensity to allow the identification of differences in growth capabilities and/or acclimation mechanisms when organisms are exposed to planetary simulations. Adding factors like a different spectrum and the anoxic atmosphere could make a saturating light become a stress factor and cells could incur photoinhibition.

To assess the most suitable light intensity, cultures at a starting OD_750_ of 0.2 were grown in a wide range of intensities, namely 10, 20, 40, 60, 80, 100 and 150 µmol of photons m^-2^s^-1^, maintaining the temperature at 25°C. Growth was followed by monitoring OD_750_, and, from OD_750_ values, the maximum growth rate µ was calculated (Figure 3). The growth rate was increased with increasing light intensities, up to 60 µmol of photons m^-2^s^-1^, where final OD_750_ reached the highest value of all conditions. With higher intensities, namely 80, 100 and 150 µmol of photons m^-2^s^-1^, growth was instead reduced, reaching low final OD_750_ values even though the maximum growth rates, registered in the first two days, was not significantly different from the ones exhibited at 10 and 20 µmol of photons m^-2^s^-1^. This growth rate was however not sustained over time in cells exposed to 80, 100 and 150 µmol of photons m^-2^s^-1^.

**Figure 3.**
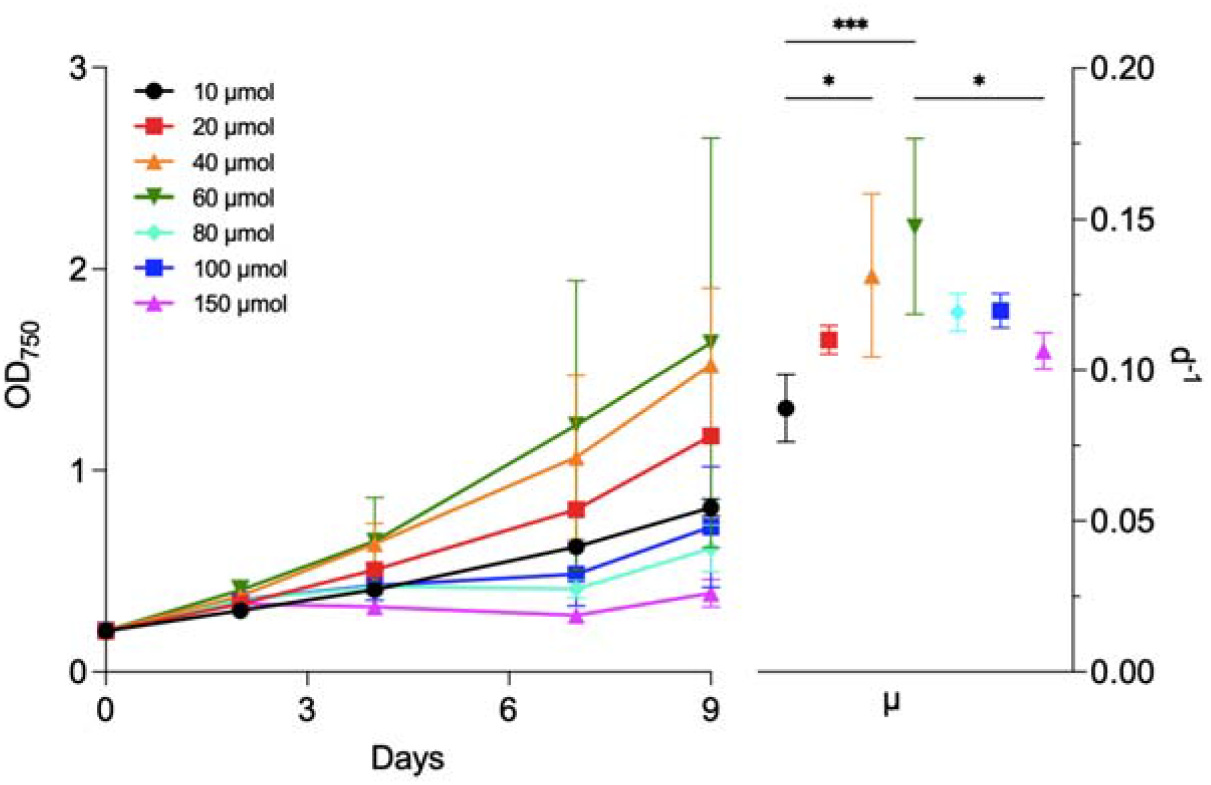
Growth of *Acaryochloris marina* sp. str. Moss Beach at different light intensities, in terms of values of OD_750_ and of growth rate (µ, d^-1^). Cells were grown at 10 (black, round), 20 (red, square), 40 (orange, triangle), 60 (green, inverted triangle), 80 (cyan, rhombus), 100 (blue, square) and 150 (pink, triangle) µmol of photons m^-2^s^-1^. OD_750_ was registered every 2-3 days (left), and, from OD_750_ values, the maximum growth rate (µ) for each condition was calculated (right). Data are presented as the mean of at least four biological replicates, for values of µ one-way ANOVA was carried out, ***: p-val <0.0005, *: p-val <0.05.

As for the previous experiment, diagnostic evaluation of the photosynthetic apparatus and pigment content was carried out on cells grown at the tested light intensities. *In vivo* absorption and low temperature fluorescence suggested no intensity had a major impact on cells – no marked differences were registered for any of the condition (Figure 4a, 4b). The highest mean values of chlorophyll *a* and *d* content were registered at 10 and 20 µmol of photons m^-2^s^-1^, a progressive decline of mean chlorophyll concentrations was observed at stronger intensities, starting from 40 µmol of photons m^-2^s^-1^ and becoming statistically significant at 80, 100 and 150 µmol of photons m^-2^s^-1^. Nonetheless, the percentage of chlorophyll *d* on the total chlorophyll content remained stable, around 94%, under all light intensities, except for cells exposed to 150 µmol photons m^-2^s^-1^, which showed a significant decrease (91.66%) (Figure 4c). Relative carotenoid content was evaluated by integrating peaks area of HPLC chromatograms (Figure 4d, S2). In cells exposed to 10, 20, 40 and 60 µmol of photons m^-2^s^-1^ the percentage of carotenoids on the total pigment extract was in the range 27-35%, while in cultures exposed to 80, 100, and 150 µmol of photons m^-2^s^-1^, the carotenoid percentage increased up to respectively 57.40%, 62.59% and 76.02%.

**Figure 4.**
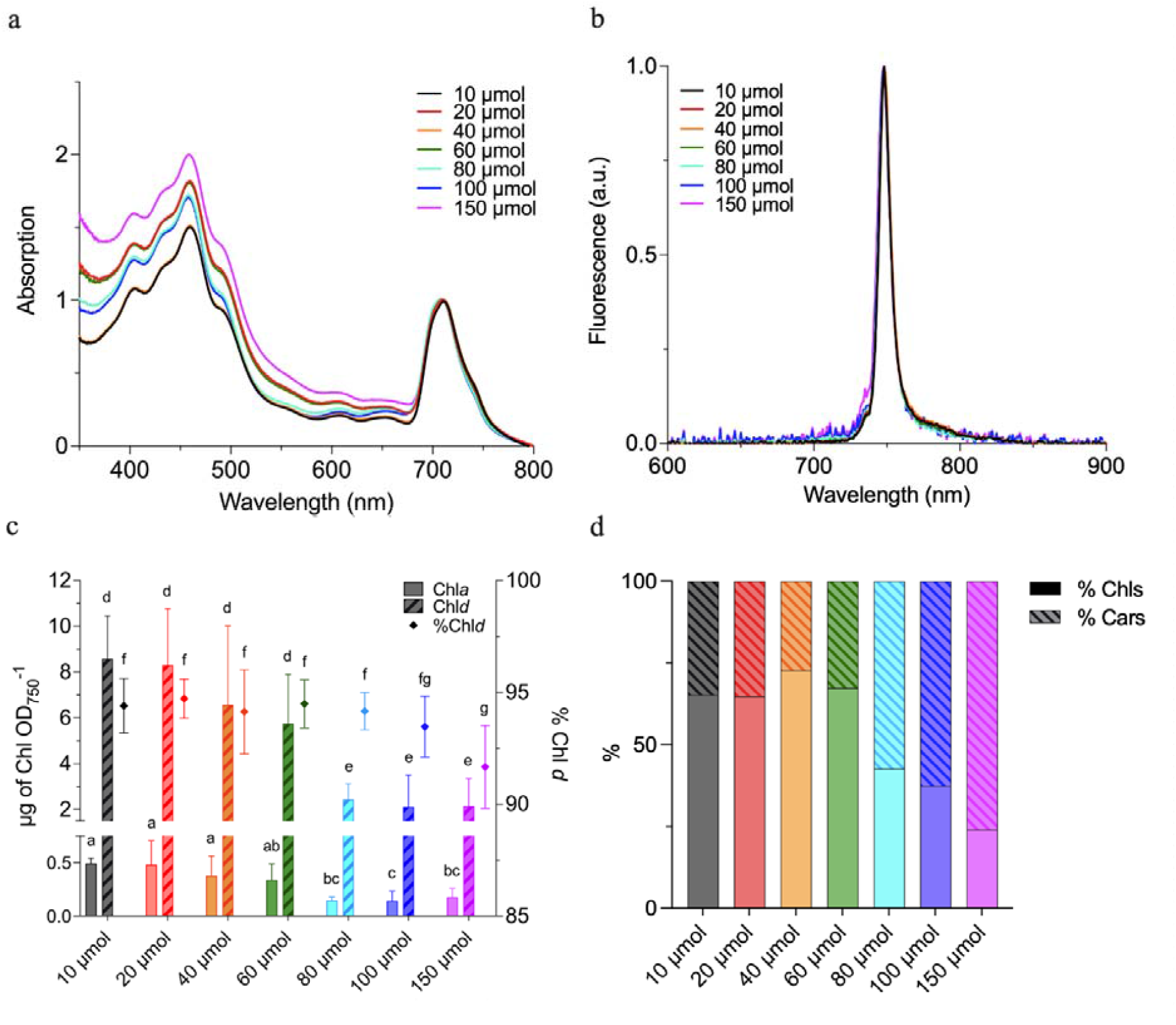
Pigment composition and photosynthetic apparatus organization in *Acaryochloris marina* sp. str. Moss Beach grown at different light intensities, 10 (black), 20 (red), 40 (orange), 60 (green), 80 (cyan), 100 (blue) and 150 (pink) µmol of photons m^-2^s^-1^. a) *in vivo* absorbance spectra of cells normalized at 710 nm, b) low temperature (77K) fluorescence emission spectra of cells, normalized at the maximum peak (748 nm), c) chlorophyll *a* (full pattern) and *d* (stripe pattern) content expressed as µg of pigment per OD_750_ and chlorophyll *d* percentage on total chlorophyll content (rhombus); statistical significance was checked with one-way ANOVA on at least four biological replicates, d) percentage of total chlorophylls (full pattern) and total carotenoids (stripe pattern) in cells extracts. Chl *a*: chlorophyll *a*; Chl *d*: chlorophyll *d*.

Even though 40 and 60 µmol of photons m^-2^s^-1^ promote the highest growth rates, these intensities result nearly saturating as shown by the initial decline of mean chlorophyll content, which is exacerbated with increasing intensities. Further experiments were hence performed at 20 µmol of photons m^-2^s^-1^.

### 3.2 Planetary simulation: M-dwarf starlight and primeval atmosphere

Different spectral qualities of light can trigger, in cyanobacteria, acclimations involving a remodeling of the photosynthetic apparatus. In order to assess whether this was the case for *A. marina* sp. str. Moss Beach under the simulated M-dwarf spectrum, cells were transferred from white light to the simulated M-dwarf starlight (M7), and, for comparison, to a simulated Solar starlight (SOL) and to a monochromatic far-red light (FR). Growth was followed for 21 days, a timing that canonically allows for light acclimations to be completely achieved. An initial lag phase of 6 days in growth was registered in all light conditions, after which cells started to grow. Values of OD_750_ revealed a noticeable growth in both FR and M7 light, with a maximum rate respectively of 0.19 d^-1^ and 0.18 d^-1^. On the other hand, cells exposed to SOL light had a lower final OD_750_ as well as a lower maximum growth rate (0.15 d^-1^) (Figure 5).

**Figure 5.**
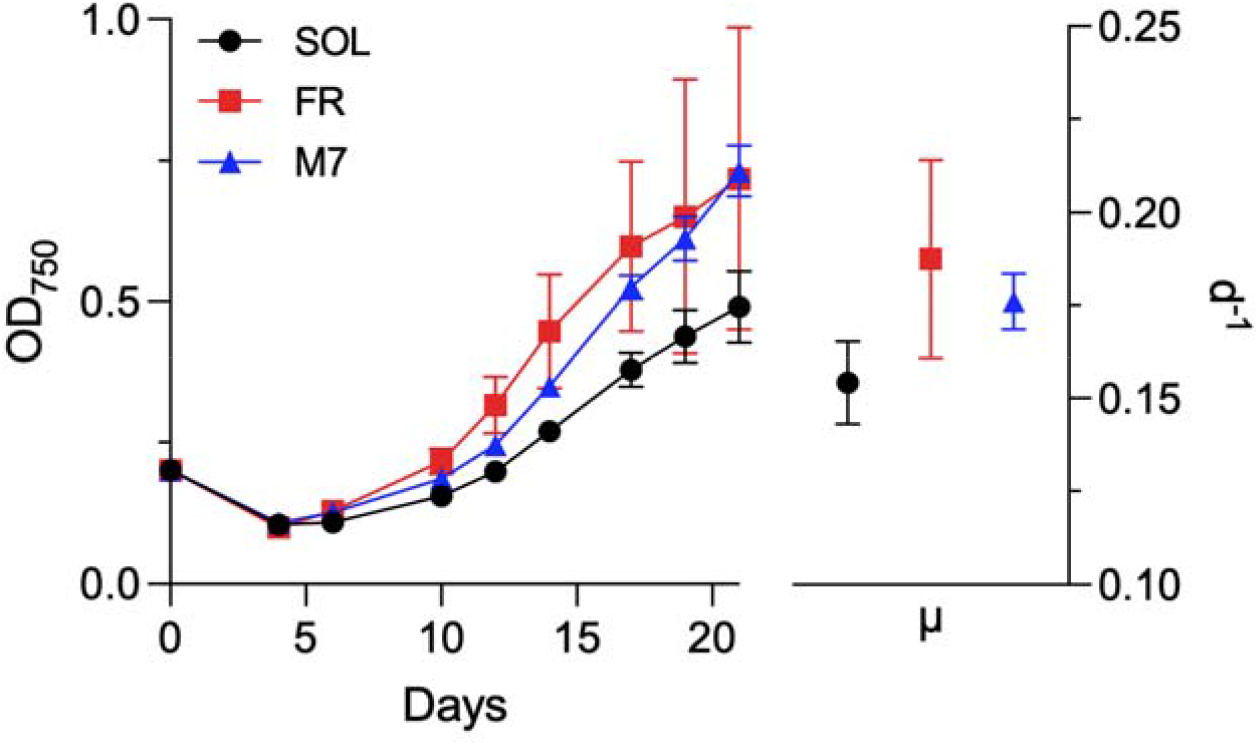
Growth of *A. marina* sp. str. Moss Beach during acclimation to the simulated stellar spectra. Growth of cells exposed to solar (SOL, black), far-red (FR, red) and M-dwarf (M7, blue) light was monitored with measures of OD_750_ every 2-4 days (left), from these values, the maximum growth rate (µ) during the 21 days was measured (right). Data are presented as the mean of three biological replicates. SOL: simulated solar light; FR: far-red light; M7: simulated M-dwarf starlight.

During acclimation to the different spectra, *in vivo* absorbance of cells was registered periodically, with no significant change of absorption spectra: after 21 days, there was no alteration from cell absorption spectrum at day zero, in any of the conditions (Figure 6a). Comparably, chlorophylls content as well as the percentage of chlorophyll *d* on the total chlorophyll content remained stable in all the tested conditions (Figure 6b). Low temperature fluorescence coherently indicated that no modification involving the emission ranges of the photosynthetic apparatus were activated under M-dwarf light, nor in any of the other light regimes: emission spectra were analogous, with the main peak being at 748 nm in all of the conditions (Figure 6c). Finally, cells were imaged with confocal microscopy to evaluate eventual changes in morphology or in the localization of chlorophylls (Figure 6d). Micrographs revealed similar shape and dimension of cells acclimated to all tested light, as well as comparable spatial distribution of chlorophyll *a* and *d* autofluorescence.

**Figure 6.**
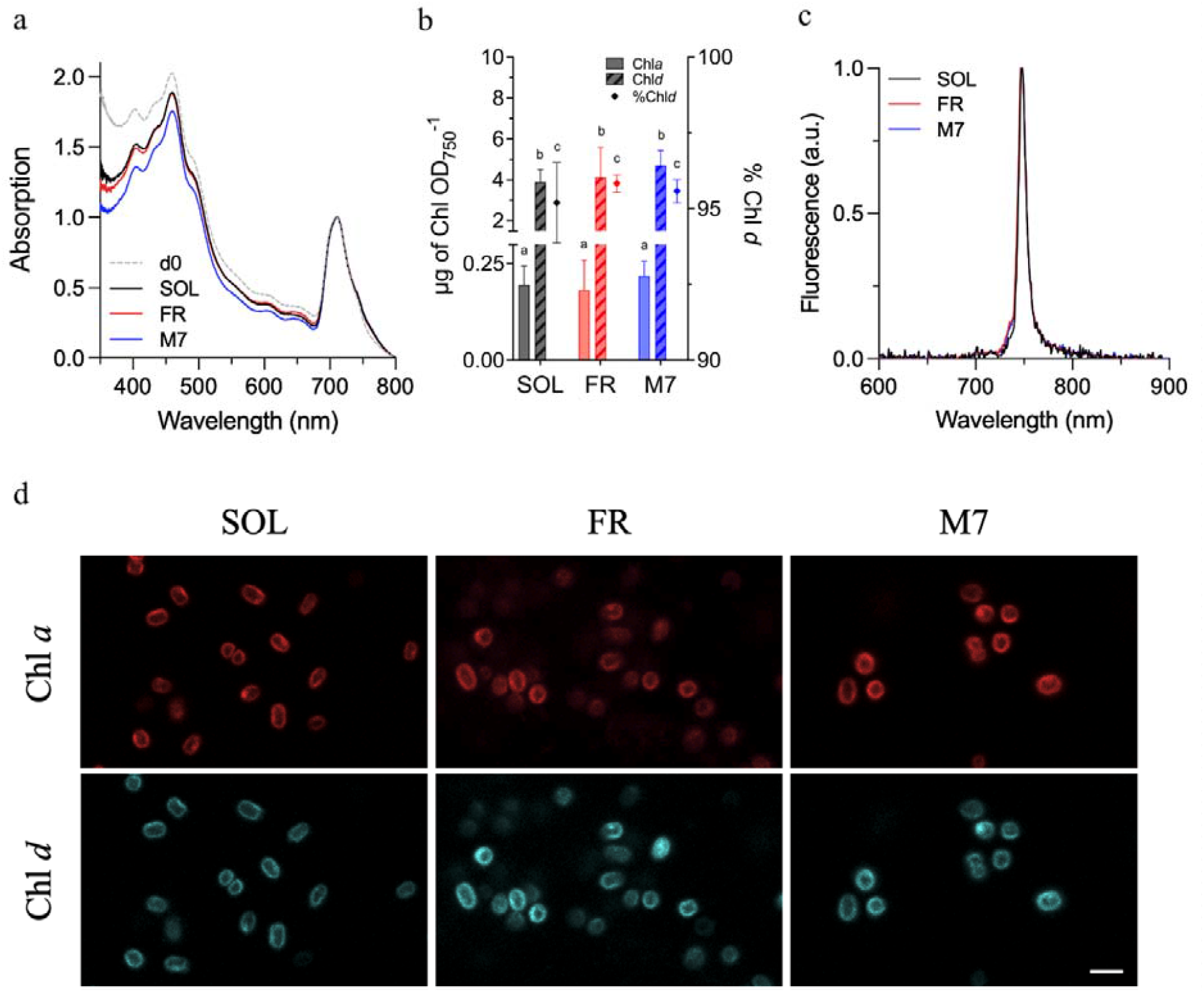
*A. marina* sp. str. Moss Beach features after acclimation to SOL, M7, FR light. a) *in vivo* absorption spectra of cell before (day zero, d0) and after acclimation; b) content of chlorophyll *a* (full pattern) and *d* (stripe pattern) in terms of µg of pigment per OD_750_, percentage of chlorophyll *d* in cell extracts; statistical significance was assessed with one-way ANOVA on three biological replicates; c) low temperature (77K) fluorescence emission spectra of cells; d) representative confocal images of respectively SOL-, FR- and M7-acclimated cells, red indicates the autofluorescence of chlorophyll *a*, cyan indicates the autofluorescence of chlorophyll *d*, scale bar corresponds to 2.5 µm. d0: day zero; SOL: simulated solar light; FR: far-red light; M7: simulated M-dwarf starlight; Chl *a*: chlorophyll *a*; Chl *d*: chlorophyll *d*.

*A. marina* sp. str. Moss Beach cultures acclimated to SOL, FR, and M7 light were used to evaluate the response and performances in simulated planetary conditions comprehensive of a modified atmosphere. Liquid cultures of OD_750_ 0.55 were placed in the atmosphere simulating chambers, ASCs, under the respective light source (SOL, FR, and M7) for 72 hours. The initial atmosphere inside the ASCs was 95% N_2_ and 5% CO_2_, reproducing a primeval anoxic atmosphere, similar to the atmosphere of Archean Earth (Catling & Zahnle, 2020; Kaltenegger et al., 2020).

After 72 h of exposure, strong growth was registered in all conditions (Figure 7a), indicating that the initial anoxic atmosphere was not preventing cell survival and reproduction. Interestingly, the simulated M-dwarf (M7) promoted the strongest growth, with cells in SOL and FR having a lower final OD_750_.

**Figure 7.**
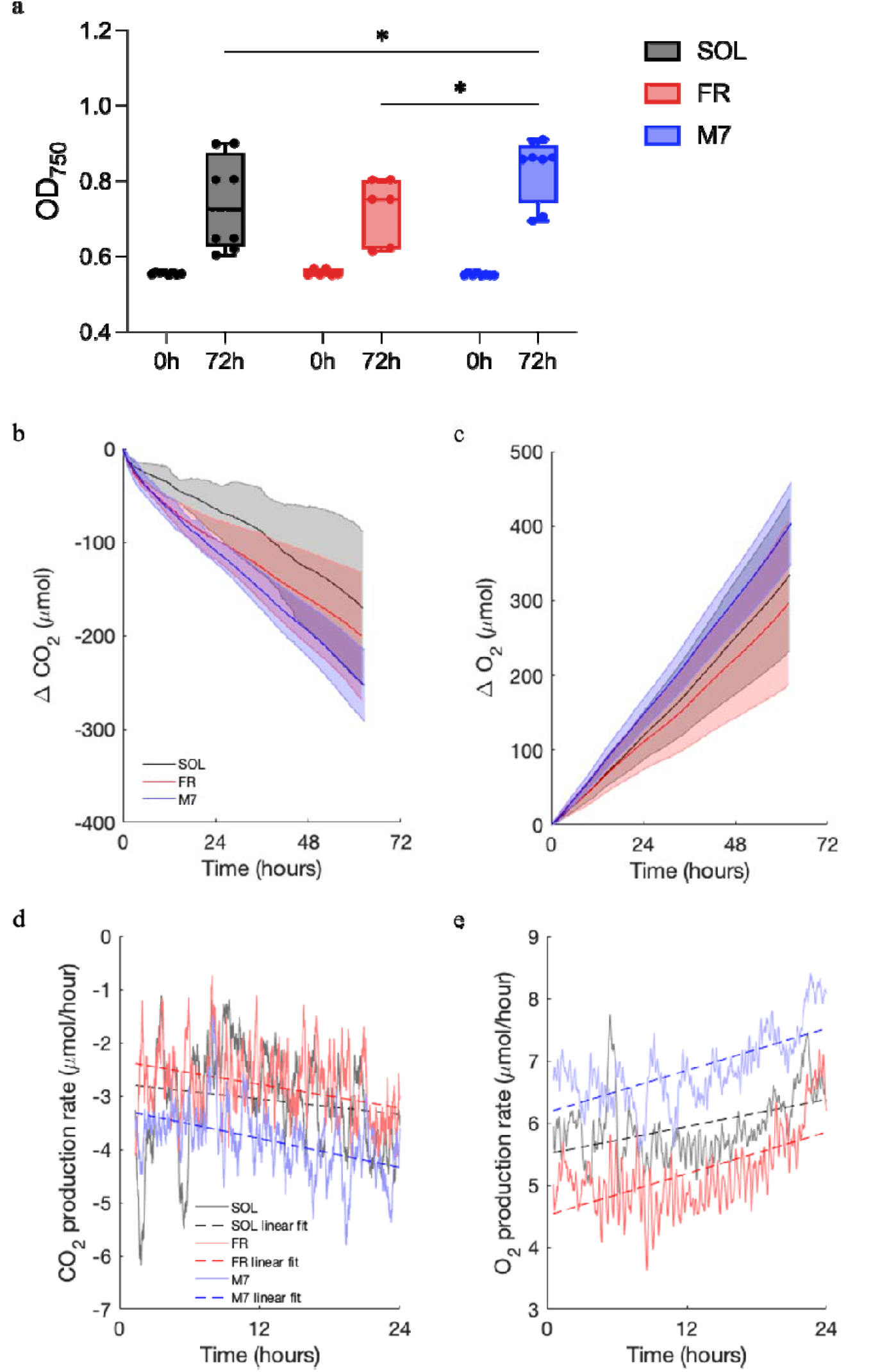
Growth and gas consumption inside the ASCs under SOL, FR and M7 light. a) initial (0h) and final (72h) OD_750_ values, b) variation of CO_2_ concentration, c) variation of O_2_ concentration, d) CO_2_ production rate, e) O_2_ production rate. For b) and c) T_0_ was set at 5 h to exclude the initial period of gas equilibration inside the ASCs. For d) and e) the last 24 hours of exposure were considered, to exclude the initial acclimation of organisms to the anoxic/microoxic environment. SOL: simulated solar light; FR: far-red light; M7: simulated M-dwarf starlight; CO_2_: carbon dioxide; O_2_: oxygen.

Coherently, cells in M7 showed a higher capability of CO_2_ consumption and O_2_ release, with final values of ΔCO_2_ and ΔO_2_ being respectively the lowest and the highest registered with respect to SOL and FR (Figure 7b, c). As highlighted in Figure 7d, e, under M7 and FR light the production rates of O_2_ and CO_2_ respectively increase and decrease faster than in SOL light, where indeed the lowest final OD_750_ value was encountered.

To assess the surface biosignature that an organism with the features of *A. marina* sp. str. Moss Beach could generate, the reflectance spectrum of cells was recorded, presented in Figure 8 along with the reflectance spectra of two other cyanobacterial strains previously tested under the starlight simulator, *Synechocystis* sp. PCC6803 and *Chlorogleopsis fritschii* PCC6912 (Battistuzzi et al., 2023a; 2023c). Reflectance spectra were recorded on cultures acclimated to M7 light, *in vivo* absorption of *S*. sp. PCC6803 and *C. fritschii* PCC6912 acclimated to SOL and M7 light are available in Figure S3.

**Figure 8.**
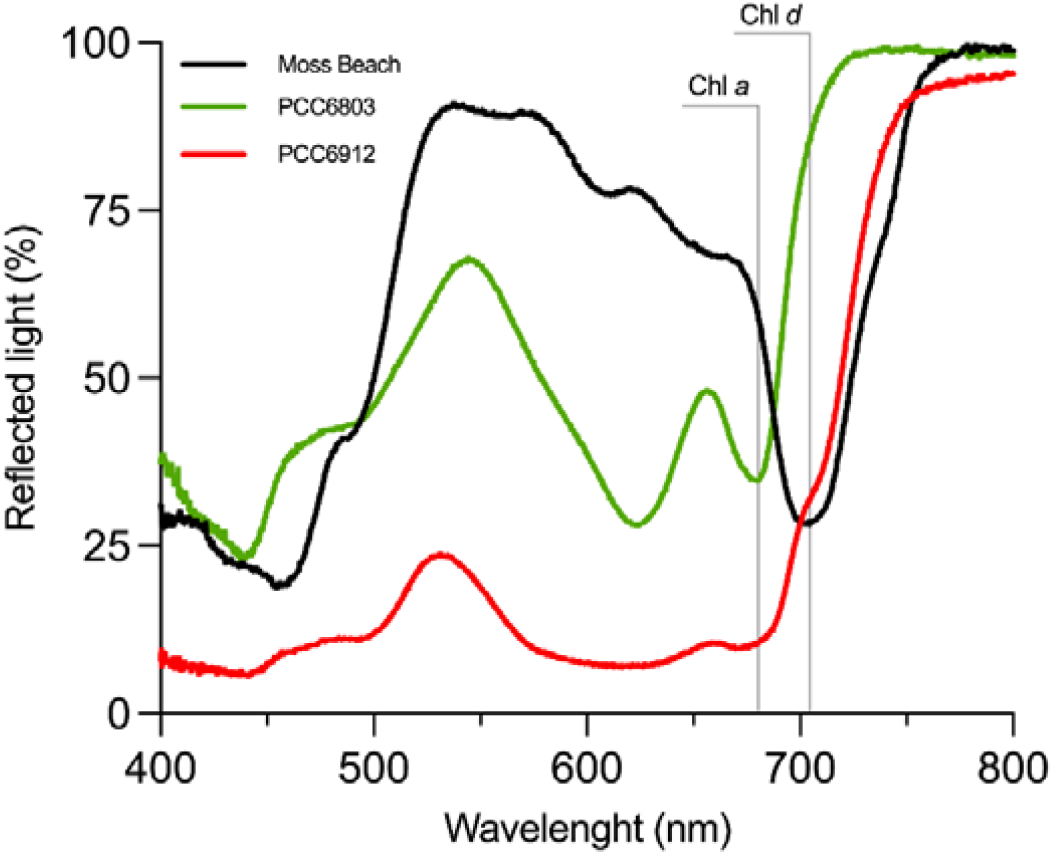
*In vivo* reflectance spectra of liquid cultures of *A. marina* sp. str. Moss Beach (black), *Synechocystis* sp. PCC6803 (green) and *Chlorogleopsis fritschii* PCC6912 acclimated to the M7 light recorded with a Flame spectrometer (OceanOptics). Moss Beach: *A. marina* sp. str. Moss Beach; 6803: *Synechococcus* sp. PCC6803; 6912: *Chlorogleopsis fritschii* PCC6912; Chl *a*: chlorophyll *a*; Chl *d*: chlorophyll *d*.

The Vegetation Red-Edge is usually found in the reflectance spectra of Earth surface portions covered by oxygenic photosynthetic organisms. As noticeable from the spectrum of *S*. sp. PCC6803, the VRE consists in a steep increase in reflectance between 680 nm and 750 nm originating from the contrast of the high absorption capacity of Chl *a* in the red and the scattering of infrared light due to cellular structures. This feature resulted altered in *C. fritschii* PCC6912 cells acclimated to M7 light, where a shift in reflectance was present above 700 nm, likely ascribable to the presence of small amounts of chlorophyll *d* and *f. In vivo* absorption of cultures used to obtain the reflectance spectra highlight indeed a shoulder of absorption beyond 700 nm in *C. fritschii* PCC6912, indicating the occurrence of the FaRLiP response in this strain under M7 irradiation (Figure S3, Battistuzzi et al., 2023a). In *A. marina* sp. str. Moss Beach, due to the abundance of Chl *d*, the steep rise in reflectance was instead completely shifted toward longer wavelengths, between 710 nm and 780 nm, in what could be referred to as a *Chl* d*-near-infra-red edge*, Chl *d-*NIR edge.

## 4. Discussion

Some of the planned space mission from NASA and ESA are aimed at characterizing the atmospheres of exoplanets, also in the search for biosignatures. In particular, HWO (NASA) will be able to detect the presence of O_2_, that astrobiologists indicate as one of the strongest biosignatures to search for. As on Earth the accumulation of O_2_ in the atmosphere is due to oxygenic photosynthesis, operated by cyanobacteria, algae and plants, for years the community has been debating on the feasibility and efficiency of this metabolism on exoplanets. In this work we focused on this possibility to occur around M-dwarf stars, as they are considered good targets for their long lifespan and the abundancy of rocky exoplanets identified as habitable harbored in their stellar systems (Gale & Wandel, 2016; Shields et al., 2016). The challenge for oxygenic photosynthesis to have evolved with an M-dwarf spectrum resides in the long wavelength limit of this metabolism. On Earth, oxygenic photosynthetic organisms rely on visible light, and can only marginally harvest far-red light, while M-dwarfs emission spectra offer very low visible and high far-red and infra-red irradiance (Chabrier et al., 1996).

Laboratory simulation have however proven how an M-dwarf spectrum does support strong growth in cyanobacteria and microalgae, and moderate growth in non-vascular plants that evolved to harvest visible light (Battistuzzi et al., 2023a; 2023b). Nonetheless, this doesn’t imply that analogous organisms should be expected to have evolved under such spectrum: a red-shifted irradiance might have selected “red-shifted adaptations” as for instance modified antennae (Chitnavis et al., 2024) or *two color* reaction centers absorbing respectively visible and near-infra-red light. (Takizawa et al., 2017).

Considering the red-shifted adaptations of oxygenic photosynthesis that have evolved on Earth, in this work we tested the Moss Beach strain of *Acaryochloris marina*, a unique species with chl *d*-bearing reaction centers and permanent red-shifted absorption, to assess whether its photosynthetic apparatus is a good fit under M-dwarf-like irradiance, also in terms of O_2_ production rates.

Due to the divergence of their light harvesting systems and chlorophyll environment, *A. marina* strains exhibit a high variability in the fluorescence properties of chl *d* and in the spectral features. Ulrich and colleagues (2024) identified three main spectral types depending on fluorescence emission features, which reflect differences in chl energy levels and in chl-binding proteins. From the *intermediate-wavelength* spectral type, with emission maximum at 727-731 nm, diverged both the *short-wavelength* and the *long-wavelength* spectral types, with emission maxima respectively at 721-724 nm and 738-748 nm. The analysis of low temperature fluorescence allowed to understand the spectral type of the strain Moss Beach, that, presenting a fluoresce peak at 748 nm, should be included in the *longer wavelength* group, to which also its close relative *A. marina* S15 belongs (Kiang et al., 2022; Ulrich et al., 2024). This locates *A. marina* sp. str. Moss Beach in the group of oxygenic photosynthetic organisms with the strongest red-shifted constitutive emission so far discovered, making it an excellent target to test Earth “red-shifted adaptions” in M-dwarf simulated light.

Results of the preliminary acclimation to the simulated stellar spectra, M7, reproducing an M-dwarf star, highlight how, differently from other species previously tested, *A. marina* sp. str. Moss Beach does not need to modulate the pigment content nor its photosynthetic apparatus in order to efficiently grow with the M7 light. When tested in the same set up, cyanobacteria like *Chlorogleopsis fritschii* sp. PCC6912 and *Synechococcus* sp. PCC7335, underwent severe remodulations of the photosynthetic apparatus, aimed at harvesting the abundant red and far-red light available. *C. fritschii* PCC6912 exhibited the FaRLiP response (Battistuzzi et al., 2023a), maximizing far-red light absorption, while *S*. PCC7335 strongly modulated the content of accessory pigments via chromatic acclimation type 3, maximizing red light absorption (Liistro et al., 2024). The unaltered absorption features and pigment content observed for *A. marina* sp. str. Moss Beach underlined, instead, how this strain is already well suited to photosynthesize under the simulated M7 stellar spectrum, growing even better than when exposed to the simulated solar light (SOL). Interestingly, during acclimation growth rates were higher than what exhibited in previous experiments conducted in white light: SOL, FR and M7 lights all provide a portion of far-red photons, in variable amounts, which appears to be crucial for this strain to drive faster growth. Indeed, at the end of the acclimation experiment performed in terrestrial atmosphere, the highest OD_750_ were observed in M7 light and monochromatic FR light, while in SOL light, which provided the shortest amount of far-red photons of all three tested lights, the final OD_750_ was the lowest. Studies conducted on *A. marina* sp. str. MBIC11017, found similar results: for this strain as well, a stronger growth was registered when cells were exposed to far-red light rather than visible light. This was paired with a stronger photosynthetic rates, indicating longer wavelength drive photosynthesis more efficiently than visible ones (Behrendt et al., 2012).

Considering all the conditions tested in this study, *A. marina* sp. str. Moss Beach showed a maximal growth rate reaching 0.19 d^-1^, comparable to the one registered in the reference strain of *A. marina*, MBIC11017 (Swingley et al., 2005).

With *A. marina* sp. str. Moss Beach acclimated to SOL, FR and M7 spectra, we then proceeded with planetary simulation, combining the light treatment with a simulated planetary primeval atmosphere non-oxygenated (95% N_2_ and 5% CO_2_). Cells grew consistently in all the light conditions, suggesting the initial absence of oxygen had no major detrimental effects. Most importantly, the M7 starlight promoted the highest growth when combined with the anoxic atmosphere, with respect with SOL and FR light. Cells were indeed efficiently sequestrating CO_2_ and producing O_2_, having the strongest rates under the simulated starlight when compared with cells in SOL and FR light.

Nonetheless, the O_2_ productivity in *A. marina* sp. str. Moss Beach is noticeable if compared with the strains previously tested in this set-up. After 72h of exposure to simulated planetary conditions (starlight spectrum coupled with primeval atmosphere), we registered the production of approximately 400 µmol of O_2_. *C. fritschii* PCC6912 and *S*. sp. PCC6803 reached similar O_2_ production after 48 hours under the same spectrum and in the same atmosphere tested in this work (Battistuzzi et al., 2023c), which is interesting considering they are both fast-growing strains and they were exposed to a higher total irradiance (30 µmol of photons m^-2^s^-1^). *S*. sp. PCC7335, a relatively slow-growing organism, took 7 days to produce the same amount under the same spectrum – however, it must be considered that the total irradiance of the experiment was set to 10 µmol of photons m^-2^s^-1^, half the one used in this study (Liistro et al., 2024). If photosynthetic metabolisms like the one selected in *A. marina* sp. str. Moss Beach have evolved outside our Solar System, our data support the potentiality of the generation of atmospheric biosignature via the release of O_2_. Nevertheless, the planetary features and the eventual vegetation coverage would definitively have an impact on the potential accumulation of this gas, hence on the possibility of its detection.

Moreover, the reflectance spectrum of *A. marina* sp. str. Moss Beach cells underlines how, if such metabolism is present on exoplanets orbiting M-dwarf stars, its fingerprint would be highly characteristic, with a VRE profoundly shifted toward longer wavelength, becoming a *near-infra-red-edge*, NIRE.

Overall, *A. marina* sp. str. Moss Beach is an excellent example of how evolution on Earth has led oxygenic photosynthesis to red-shifted adaptations that efficiently photosynthesize with an M-dwarf like spectrum, underlining the possibility of natural selection to push highly functional metabolisms forward their energetic limits, as in the case of *A. marina* species.

## 5. Conclusions

The *A. marina* species constitute fine evidence of how evolutive pressure can lead oxygenic photosynthesis not only toward visible-harvesting forms but also toward constitutive red-shifted forms.

In this work, the Moss Beach strain of *A. marina* was tested for the first time under a simulated M-dwarf light, evaluating also its capacity of producing O_2_ in an anoxic atmosphere enriched in CO_2_. The results of this work highlight how the photosynthetic apparatus of *A. marina* sp. str. Moss Beach represents a very good, constitutively red-shifted, fit for oxygenic photosynthesis under M-dwarf stars, leading to the release of considerable amounts of O_2_ with a strong impact on the initial anoxic atmosphere. Surprisingly the O_2_ production rate resulted comparable to the visible-light user cyanobacteria that have been previously tested. If evolved on exoplanets orbiting around an M-dwarf, this kind of metabolism could hence generate a gaseous biosignature. The data of O_2_ release here presented are of crucial interest for exoplanet habitability models and represent a reference to interpret the data that will eventually result from future space missions like ARIEL (ESA) and HWO (NASA), that are going to investigate exoplanets’ atmospheres. Regarding the possibility of such organisms to generate a surface biosignature, the pigment content of *A. marina* sp. str. Moss Beach is unaltered under the solar or the M-dwarf spectra, resulting in a characteristic reflectance spectrum with a Vegetation Red-Edge sensibly shifted toward longer wavelengths, in a *near-infra-red edge*.

## CRediT authorship contribution statement

EL: Conceptualization, Formal analysis, Investigation, Methodology, Data curation, Visualization, Writing – original draft, Writing – review and editing; BB: Formal analysis, Data curation, Writing – review and editing; MNP: Resources, Conceptualization, Writing – review and editing; NYK: Resources, Conceptualization, Writing – review and editing; NLR: Resources, Conceptualization, Investigation, Data Curation, Supervision, Writing – original draft, Writing – review and editing.

## Acknowledgments

Authors would like to thank Marianna Quadri, Mariano Battistuzzi, Lorenzo Cocola and the staff of the Imaging Facility of the Department of Biology of the University of Padova for the scientific and technical support.

## Footnotes

1 From www.exoplanets.nasa.gov consulted on April 8, 2025

