## Supplementary material for "Is constitutive red-shift an advantage for oxygenic photosynthesis under M-dwarf starlight? Insights from *Acaryochloris marina* sp. str. Moss Beach"

a b

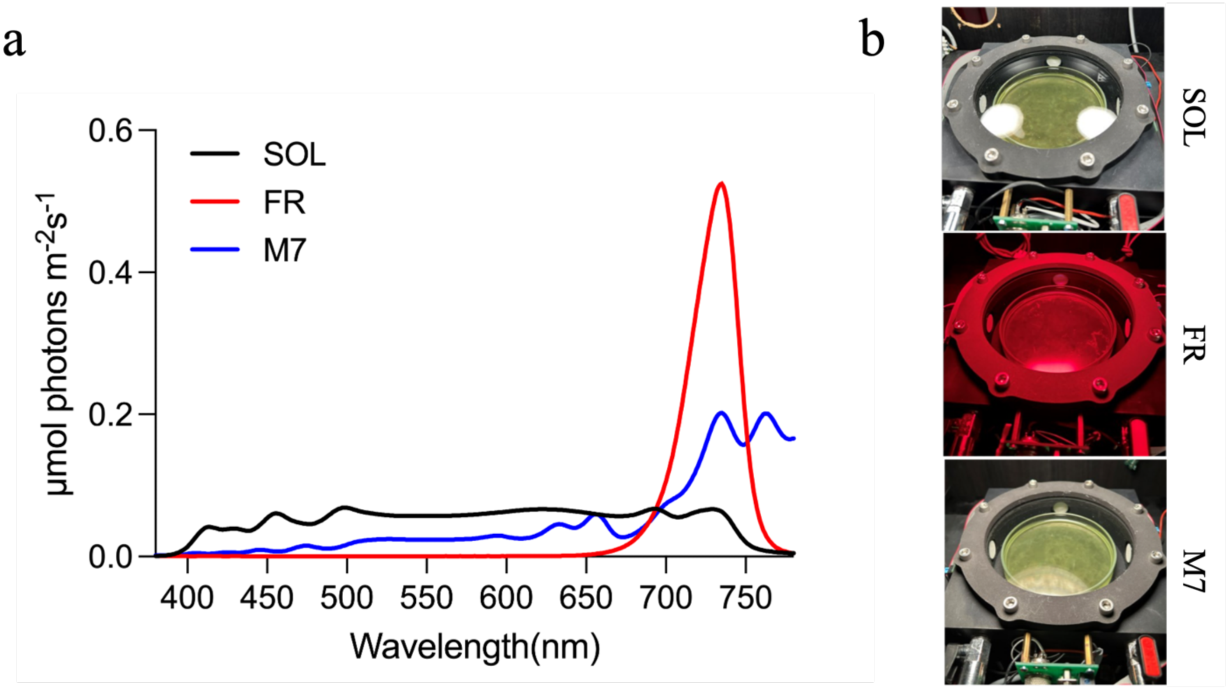


c

| | SOL | | FR | | M7 | |
|---|---|---|---|---|---|---|
| Wavebands | µmol of photons $m^{-2}s^{-1}$ | % | µmol of photons $m^{-2}s^{-1}$ | % | µmol of photons $m^{-2}s^{-1}$ | % |
| UV 380–399 nm | 0.08 | 0.4 | 0.01 | 0.05 | 0.04 | 0.2 |
| Blue 400-499 nm | 4.61 | 23.05 | 0.04 | 0.2 | 0.87 | 4.35 |
| Green 500-599 nm | 5.95 | 29.75 | 0.04 | 0.2 | 2.43 | 12.15 |
| Red 600-699 nm | 6.28 | 31.4 | 1.38 | 6.9 | 4.04 | 20.2 |
| Far-red 700-780 nm | 3.08 | 15.4 | 18.53 | 92.65 | 12.62 | 63.1 |
| total VIS 380-699 nm | 16.92 | 84.6 | 1.47 | 7.35 | 7.38 | 36.9 |
| total light 380-780 nm | 20 | 100 | 20 | 100 | 20 | 100 |

Figure S1. Light and ASCs set-up. a) solar (SOL), far-red (FR) and M-dwarf (M7) simulated spectra, , b) pictures of the organisms inside the ASCs (right), exposed to the respective spectra, c) repartition of light in terms of µmol of photons $m^{-2}s^{-1}$ and percentage in the wavebands UV, blue, green, red, far-red, total VIS light, total light recorded with a LI-800 spectrometer (LICOR, USA). SOL: solar light, M7: M-dwarf simulated light, FR: far-red light, UV: ultraviolet light, VIS: visible light.

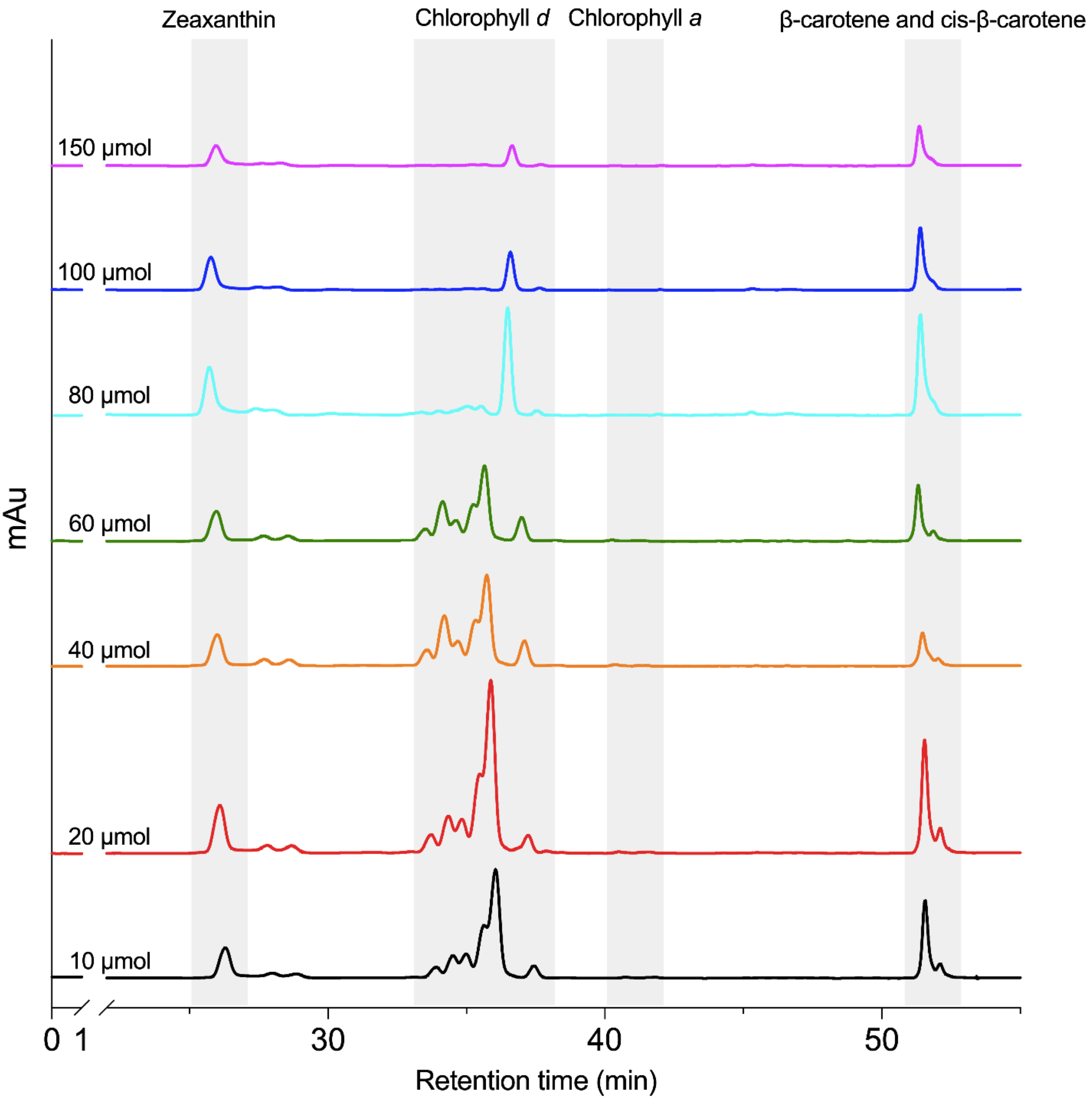


Figure S2. Chromatograms at 440 nm of 100% methanol exctracts of *A. marina* str. Moss Beach after exposure to 10 (black), 20 (red), 40 (orange), 60 (green), 80 (cyan), 100 (blue), 150 (pink) µmol of photons $m^{-2}s^{-1}$. Grey bands indicate elution of respectively zeaxanthin, chlorophyll *d*, chlorophyll *a*, β-carotene and cis-β-carotene.

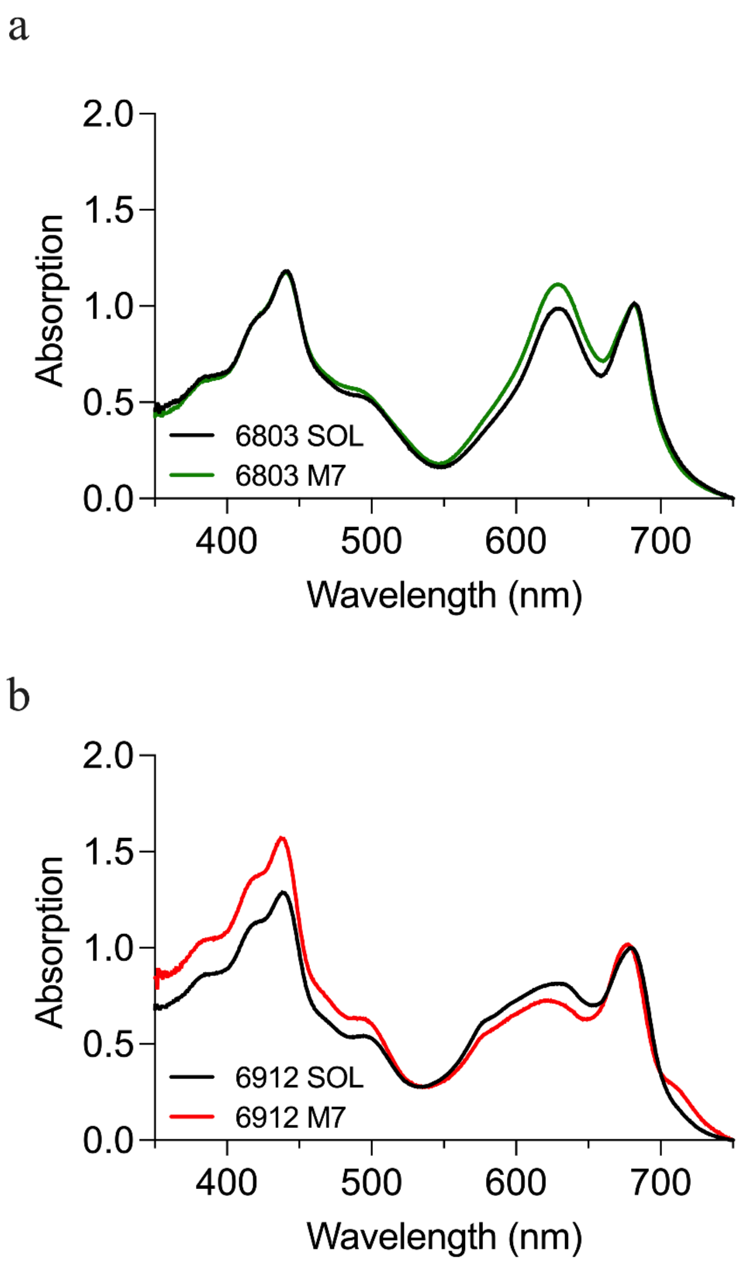


Figure S3. *In vivo* absorption spectra of (a) *Synechococcus* sp. PCC6803 acclimated to SOL (black line) and M7 (green line) light, and (b) *Chlorogleopsis fritschii* PCC6912 acclimated to SOL (black line) and M7 (red line) light. 6803: *Synechococcus* sp. PCC6803; 6912: *Chlorogleopsis fritschii* PCC6912; SOL: simulated solar light; M7: simulated M-dwarf starlight.
